# Differential roles for DNAJ isoforms in HTT-polyQ and mutant FUS aggregation modulation revealed by chaperone network screens

**DOI:** 10.1101/2021.04.08.438944

**Authors:** Kinneret Rozales, Amal Younis, Lior Kellerman, Ronit Heinrich, Shai Berlin, Reut Shalgi

## Abstract

Protein aggregation is a hallmark of many neurodegenerative diseases^1,2^. In order to cope with misfolding and aggregation, cells have evolved an elaborate network of molecular chaperones, composed of different families^3^. But while chaperoning mechanisms for different families are well established, functional and regulatory diversification within chaperone families is still largely a mystery^4,5^. Here we decided to explore chaperone functional diversity, through the lens of pathological aggregation. We revealed that different naturally-occurring isoforms of DNAJ chaperones showed differential effects on different types of aggregates. We performed a chaperone screen for modulators of two neurodegeneration-related aggregating proteins, the Huntington’s disease-related HTT-polyQ, and the ALS-related mutant FUS (mutFUS). The screen identified known modulators of HTT-polyQ aggregation^6,7^, confirming the validity of our approach. Surprisingly, modulators of mutFUS aggregation were completely different than those of HTT-polyQ. Interestingly, different naturally-occurring isoforms of DNAJ chaperones had opposing effects on HTT-polyQ vs. mutFUS aggregation. We identified a complex of the full length (FL) DNAJB14 and DNAJB12 isoforms which substantially alleviated mutFUS aggregation, in an HSP70-dependent manner. Their naturally occurring short isoforms were unable to form the complex, nor to interact with HSP70, and lost their ability to reduce mutFUS aggregation. In contrast, the short isoform of DNAJB12 significantly alleviated HTT-polyQ aggregation, while DNAJB12-FL aggravated HTT-polyQ aggregation. Finally, we demonstrated that full-length DNAJB14 ameliorated mutFUS aggregation compared to DNAJB14-short in primary neurons. Together, our data unraveled distinct molecular properties required for aggregation protection in different neurodegenerative diseases, and revealed a new layer of complexity of the chaperone network elicited by naturally occurring J-protein isoforms, highlighting functional diversity among the DNAJ family.

---

Protein aggregation is a hallmark of the many neurodegenerative diseases (NDs), such as Alzheimer’s disease, Parkinson’s disease, Huntington’s disease and amyotrophic lateral sclerosis (ALS)^1,2^. Importantly, while protein aggregation is common to many different NDs, evidence suggests different biophysical properties of aggregates formed in different diseases^8–10^.

Molecular chaperones are the master regulators of protein folding and aggregation in the cell^3^. As such, they have been the focus of various studies attempting to understand their role as potential modifiers of ND-related protein aggregation^11^. Of particular interest is the HSP70 network, including a diverse family of HSP40 co-chaperones (also called DNAJs), and Nucleotide Exchange Factors (NEF) co-chaperones, both of which are required for HSP70 to exert its chaperoning function^4^. Interestingly, several chaperone families have undergone expansions throughout evolution, with 13 different HSP70s, and a similar number of different NEFs encoded in the human genome. The DNAJ family of co-chaperones have undergone a major expansion with over 50 different members in humans^4^. These expansions increase the combinatorial capacity of the chaperone network, with the general notion that different DNAJs confer different client specificity to HSP70s, however, for the vast majority of them, this specificity is still unknown ^4,5^.

In the context of aggregation, several chaperones have been shown to confer aggregation protection, the majority of them in the context of the Huntington’s disease-related Huntingtin protein (HTT-polyQ). Several chaperones have been found to reduce HTT-polyQ aggregation, primarily DNAJB6 and DNAJB8^6,12–15^, and a few others^7,16,17^.

In ALS, however, and in particular for the ALS-mutated FUS (mutFUS), chaperone modifiers were much less explored. The yeast HSP104 chaperone has been highlighted as a disaggregase of the ALS aggregating proteins FUS and TDP43 in yeast^18^. Using over-expression and deletion screens in yeast, significant FUS aggregation modifiers were found to include RNA-binding proteins and proteins involved in stress granules^19^. Nevertheless, WT FUS showed similar aggregation and toxicity to mutFUS in both yeast and fly models^19,20^, underscoring the importance of exploring mutFUS aggregation modulation in a mammalian system.

Here we sought to explore the functional diversity of chaperones from different families, in order to unravel functional diversification in the context of different aggregate types.

## Chaperone network screen for modulators of HTT-polyQ aggregation

To systematically characterize the functional effects of individual chaperones on protein aggregation, we first established a functional screening system for modifiers of ND-related protein aggregation phenotype in human cells. To that end, we calibrated the PulSA method^21^, originally developed for detection and quantification of HTT-polyQ protein aggregation using flow cytometry (FACS), to be used as a readout for a chaperone screen (Fig. 1A). HEK293T cells ectopically expressing the HTT-134Q-GFP disease-related protein (specifically exon1 of the HTT protein, see Fig. S1A,B) gave rise to two cell populations, whereas cells expressing the WT version (HTT-17Q-GFP) presented a single fluorescent population, with an effective fraction of AGG+ cells of zero (Fig. S1C). We next turned to screen for potential HTT-134Q chaperone modulators in a co-expression screen, using a standardized co-expressed neutral protein, DsRed (Fig. S1D,E, see Methods). The aggregate-containing cell fraction (AGG+) resulting from the co-expression of each chaperone together with HTT-134Q-GFP was normalized to a set of DsRed co-expression controls, and the log2 fold change in AGG+ fraction compared to the control was denoted as the “Aggregation modulation score” (Fig. 1A). Many biological replicates of the control were used to generate a 95% confidence interval (see Methods), allowing us to assess whether a chaperone had a significant effect on protein aggregation. Chaperones with a negative Aggregation modulation score, i.e. below the confidence interval, are those that provided a significant alleviation of the aggregation phenotype, whereas chaperones with a positive Aggregation modulation score above the confidence interval, significantly aggravated protein aggregation (Fig. 1A).

**Figure 1:**
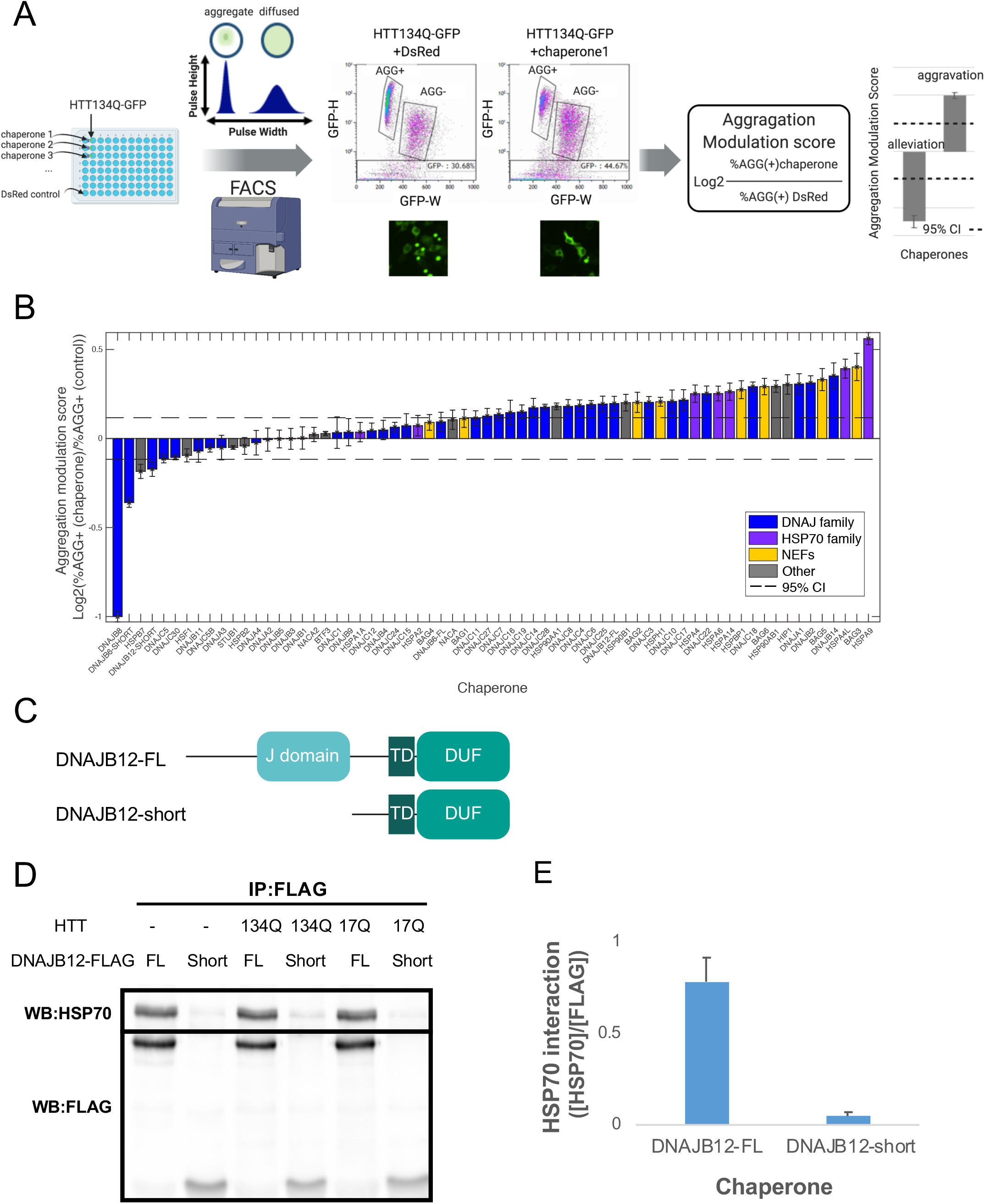
DNAJB12 isoforms with differential effects on HTT-polyQ aggregation revealed in a chaperone aggregation modulation screen. (A) Schematic cartoon of the screening framework for chaperone modulators of HTT-polyQ aggregation. First, HTT134Q-GFP was expressed in HEK293T cells together with each chaperone (Table S1), or with DsRed control. Then, samples were subject to FACS analysis using the PulSA assay^21^, considering the fluorescent peak height and width, and the fraction of aggregate-containing cells was quantified (AGG+ population). An aggregation modulation score was then calculated for each chaperone, and a 95% confidence interval (95% CI, dashed horizontal lines) was calculated according to the variation between all DsRed control replicates (N=57, corresponding to +/−2*STD). A negative Aggregation Modulation Score denotes aggregation alleviation, while a positive score denotes aggregation aggravation. (B) Aggregation modulation scores calculated for 66 chaperones showed many modulators that significantly aggravated HTT-134Q-GFP aggregation, while four chaperones significantly protected from HTT-134Q-GFP aggregation. Bars represent mean and standard errors of N = 4 independent biological replicates for each chaperone. 95% confidence intervals (dashed lines) were calculated based on 57 DsRed replicates (CI=+/−0.1168, corresponding to +/−2*STD). Chaperone families are color coded. (C) Schematic diagram of DNAJB12 isoforms. The J domain is absent in the DNAJB12-Short. TD - Transmembrane domain, DUF - Domain of unknown function. (D-E) Co-immunoprecipitation (co-IP) showed the interactions of DNAJB12 isoforms with HSP70. HEK293T cells were transfected DNAJB12-FL or -short isoforms tagged with FLAG, either alone or in the presence of either HTT-134Q-GFP or HTT-17Q-GFP. Cells were subjected to co-IP 48h later using an anti-FLAG antibody, followed by western blot with anti-HSC70/HSP70 antibody. Western with anti-FLAG antibody was perform to normalize for the amount of pulled-down chaperone. DNAJB12-FL, which contains a J-domain, interacted with HSP70 while the short isoforms lacking the J-domain showed negligible interaction. Quantification was performed using Fiji (E), mean and std of (D) is shown.

## Different DNAJB12 isoforms show differential effects on HTT-polyQ aggregation

Next, we systematically co-expressed HTT-134Q-GFP together with each of a set of 66 chaperones that were tagged with FLAG, focusing mainly on the subnetwork of HSP70 chaperones and their co-chaperones (Table S1), which have undergone a major expansion in evolution. While control cells yielded around 34% HTT-134Q-GFP aggregate-containing cells, co-expression of DNAJB8 reduced the fraction of HTT-134Q-GFP aggregate-containing cells by 50% on average, and up to ~13% (Fig. 1B,S1F). These FACS-based results were confirmed by microscopy imaging, illustrating reduced aggregate formation in DNAJB8 expressing cells (Fig. S1G, S2A-B).

Overall, our chaperone overexpression screen revealed a continuous spectrum of Aggregation modulation scores (Fig. 1B), with ~40% of the chaperones significantly aggravating HTT-134Q-GFP aggregation phenotype, and four chaperones that provided significant rescue of HTT-134Q-GFP aggregation. Importantly, the top three chaperones that we found to significantly reduce HTT-134Q aggregation, DNAJB8, DNAJB6-short and HSPB7 (Fig. 1B, S2A-C), were all previously identified using biochemical methods to provide HTT-polyQ aggregation protection in human cells^6,7,12^. Conversely, co-expression of HSP90 (the constitutive isoform HSP90AB1), aggravated HTT-134Q-GFP aggregation, with an average of 1.22 fold more aggregate-containing cells than the respective controls (Fig. 1B, S1F,G, S2A,D). This result is in agreement with a previous report on the alleviation of HTT-polyQ aggregation with HSP90 inhibitors^22^. Additionally, among the top chaperones which aggravated HTT-polyQ aggregation phenotype was DNAJA1, whose knockout was recently shown to reduce HTT-polyQ aggregation^23^. Collectively, these evidences support the validity of our screening approach.

Interestingly, some chaperones had different isoforms, denoted full-length (FL) and short, with differential effects on HTT-polyQ aggregation. Our screen found one chaperone with a previously unidentified significant rescue of HTT-polyQ aggregation (Fig. 1B): DNAJB12-short, a naturally occurring short isoform of DNAJB12. Specifically, this short DNAJB12 isoform lacks the J-domain responsible for HSP70 interactions of all J-proteins (Fig. 1C). Indeed, while the full-length isoform DNAJB12-FL strongly interacted with HSP70, the short DNAJB12 isoform did not show HSP70 interactions in co-immunoprecipitation (coIP) experiments (Fig. 1D,E). Surprisingly, DNAJB12-FL showed the opposite effect from DNAJB12-short, as it significantly elevated HTT-polyQ aggregation (Fig. 1B, S2E). Therefore, it seems that different isoforms of DNAJB12 show opposing functional effects on HTT-polyQ aggregation.

## Modulators of ALS-related mutant FUS aggregation are distinct from those of HTT-polyQ

Next, we wished to explore chaperone functional modulations of a different type of ND-related aggregate, which has not been previously explored in this context, namely the ALS-related proteins FUS^24^. We focused on a FUS mutation that occurs in patients of familial ALS, R521H^25^, localized in the Nuclear Localization Signal (NLS) region of the protein. While FUS-WT-YFP largely localized to the nucleus, the FUS-R521H-YFP mutant formed perinuclear aggregates (Fig. 2A). An additional ALS-related mutation, R518K, gave a similar pattern of localization (Fig. S3A). We then adapted our FACS-based readout to allow quantification of mutFUS aggregation, by detection of an AGG+ population in FUS-R521H-YFP expressing cells that does not appear in cells expressing FUS-WT-YFP (Fig. S3B). Here too, DsRed co-expression was chosen as a baseline control for the functional screen (Fig. S3C, see Methods).

**Figure 2:**
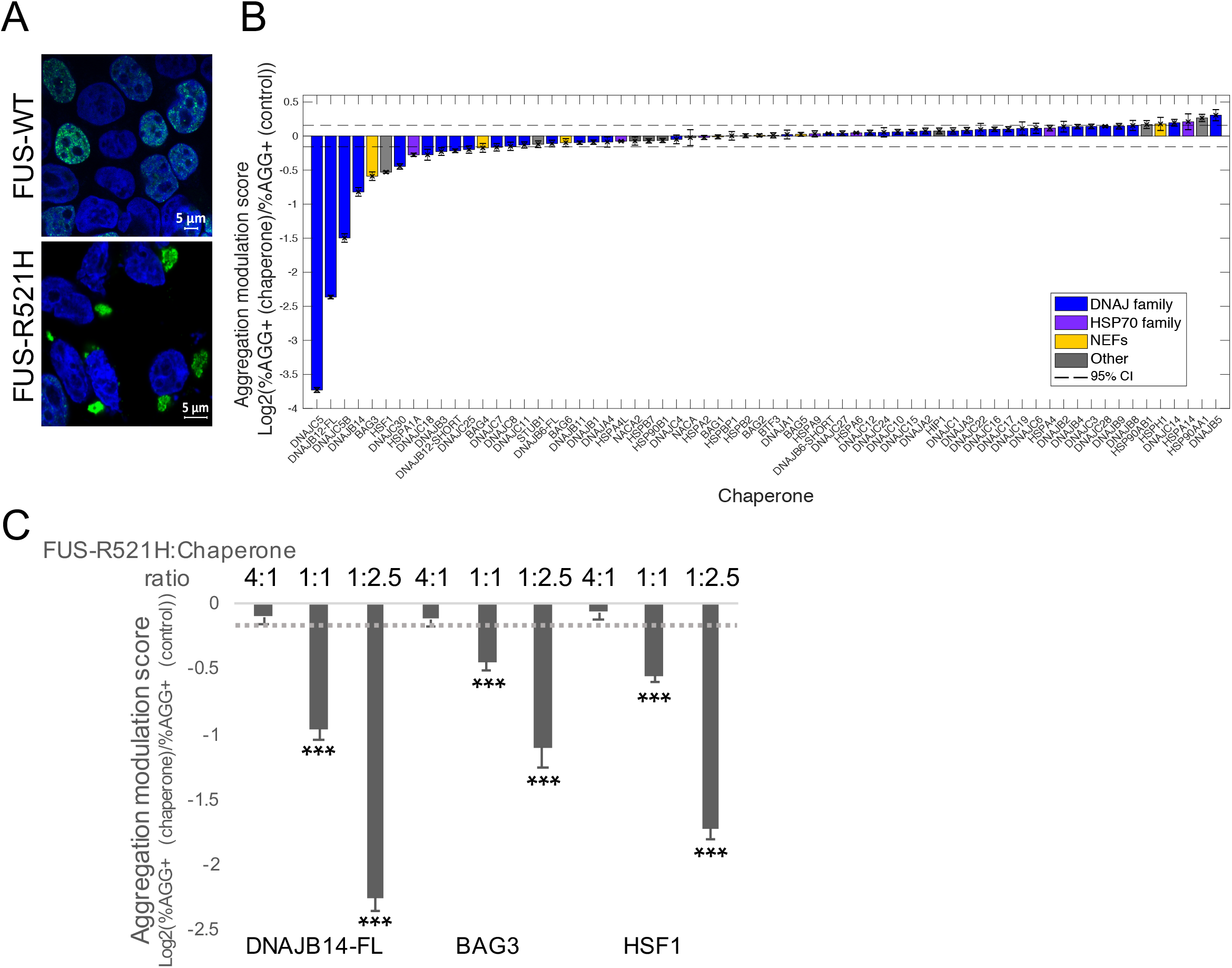
Chaperone network screen reveals modulators of mutant FUS aggregation. (A) HEK293T cells transfected with FUS-WT-YFP or FUS-R521H-YFP (green), were imaged 48h following transfection using confocal microscopy. Cells were stained with DAPI to mark nuclei (blue). Cytoplasmic perinuclear FUS-YFP signal was observed for the R521H mutants, whereas WT-FUS-YFP exhibited a predominately nuclear localization. FUS-R518K (Fig. S3A) showed a similar pattern to FUS-R521H. (B) Aggregation modulation scores for 66 chaperones revealed novel aggregation alleviators (negative scores, left side). Bars represent mean and std error of N=4 independent biological replicates. 95% confidence intervals marked with dashed lines (CI=+/−0.1577, representing 2*STD based on a total of 90 DsRed replicates). Chaperone families are color coded. (C) Aggregation modulation showed a dose dependent effect. Cells were co-transfected with different FUS-R521H-YFP:chaperone ratios, starting from high ratio of FUS DNA:chaperone DNA (2000ng/500ng respectively), equal DNA amounts as in the screen (1250ng/1250ng), and a low ratio (700ng/1800ng respectively). Results demonstrated a dose dependent rescue for all chaperones shown. Mean and std error of N=4 independent biological replicates presented. Dashed lines represent 95% CI as in (B). *** - p<0.003.

The aggregation modulation screen for FUS-R521H-YFP was then performed using co-expression of each of the 66 chaperones in our examined HSP70 network (Fig. 2B). Interestingly, while the majority of chaperones did not significantly alter FUS-R521H-YFP aggregation phenotype, two chaperones, HSP90AA1 and DNAJB5 showed a slight but significant aggravation of FUS-R521H-YFP aggregation. Furthermore, our screen revealed 11 chaperones that significantly rescued FUS-R521H-YFP aggregation, out of which seven gave a substantial rescue of more than 25% (Aggregation modulation score < −0.41, Table S1, Fig. 2B). We note that the rescue significance was robust to the FACS parameter settings (Fig. S3D). Comparison between HTT-polyQ and mutant FUS Aggregation modulation scores showed that FUS-R521H-YFP aggregation modulators were completely different than those of HTT-polyQ aggregation, and the correlation between HTT-polyQ and mutFUS Aggregation modulation scores was negligible (Fig. S3E).

We checked the extent to which these top chaperones affected cell viability. Among the top seven, three chaperone, DNAJC5, DNAJB12-FL and DNAJC5B, had a negative effect on cell viability of 15% or more (Fig. S3F). We therefore further characterized a few of the other aggregation protective chaperones. Microscopy imaging of DNAJB14, HSF1 and BAG3 co-expressing cells confirmed less aggregation of FUS-R521H-YFP compared to the control (Fig. S3G). Additionally, the same FUS-R521H-YFP aggregation rescue chaperones we identified were able to rescue the aggregation phenotype of another mutant, FUS-R518K-YFP, to a similar extent (Fig. S3H,I).

Importantly, we found FUS-R521H-YFP aggregation rescue to be dose-dependent (Fig. 2C, see Methods). For example, while the Aggregation modulation score obtained for DNAJB14 was originally −1, reflecting a 50% reduction in the fraction of FUS-R521H-YFP aggregate-expressing cells, higher doses of this chaperone promoted a 78% reduction in FUS-R521H-YFP aggregation phenotype (Fig. 2C).

## Differential roles for DNAJB14 isoforms in mutant FUS aggregation protection

DNAJB12-short, which we identified as a modulator of HTT-polyQ aggregation above (Fig. 1B), provided only marginal rescue of FUS-R521H-YFP aggregation (Fig. 2B). Interestingly, DNAJB14 also has a naturally-occurring short isoform, lacking the J-domain, the DUF and the transmembrane domain (TD) (Fig. 3A). Moreover, DNAJB14 has been previously shown to interact and co-localize with DNAJB12^26^. We next cloned the short isoform of DNAJB14 (Fig. 3A, see Methods) and tested its functional effect on FUS-R521H-YFP aggregation. In contrast to the significant rescue by DNAJB14-FL, DNAJB14-short did not rescue FUS-R521H-YFP aggregation (Fig. 3B). These trends were confirmed by image analysis of FUS-R521H-YFP aggregation in immunofluorescence images containing hundreds of cells (Fig. 3C, S4A-B, see Methods).

**Figure 3:**
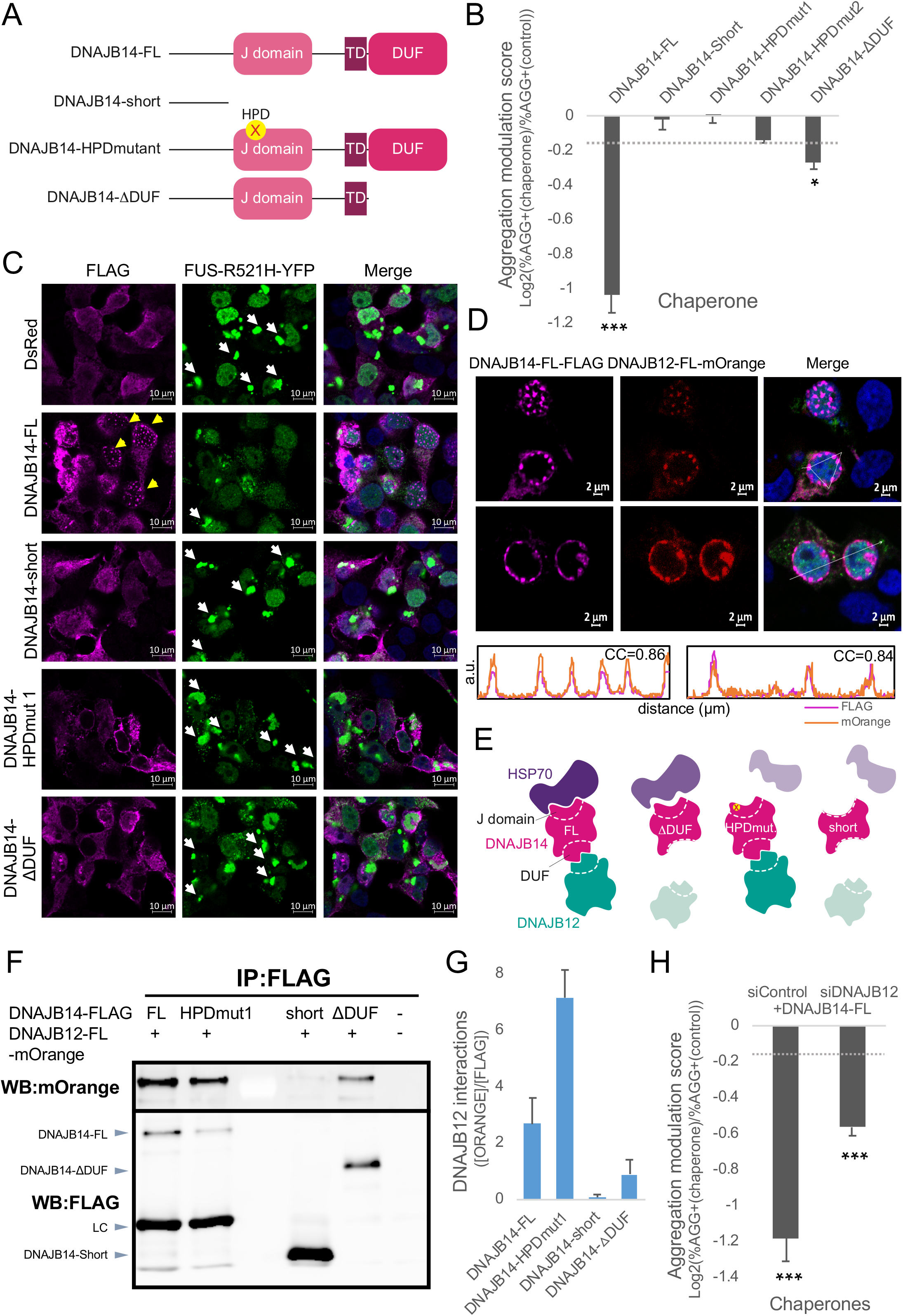
Differential roles for DNAJB14 isoforms in mutant FUS aggregation protection. (A) Schematic diagram of DNAJB14 isoforms and mutants. The J domain, TD (Transmembrane domain) and DUF (Domain of unknown function) domains are absent in the naturally-occurring DNAJB14-short isoform. HPD mutants were generated by mutating DNAJB14-FL in the HPD motif inside the J-domain, where His was changed to Gln by replacing T to G (HPDmut1) or T to A (HPDmut2). The ΔDUF mutant lacks the C-Terminal DUF domain. (B) FUS-R521H-YFP Aggregation modulation scores for DNAJB14-FL, DNAJB14-short, DNAJB14-HPDmut1 and 2, and DNAJB14-ΔDUF. Mean and std error presented for N=3 independent biological replicates. Dashed lines represent 95% CIs as in Fig. 2B. *** - p<0.003. * - p<0.05. (C) FLAG-tagged DNAJB14 isoforms and mutants co-expressed together with FUS-R521H-YFP (green). Immunofluorescence imaging (IF) was performed 48h later, using far-red anti-FLAG antibody (pink), DAPI marks nuclei (blue). Yellow arrows mark DJANGO structures, which were observed only in samples expressing DNAJB14-FL. White arrows mark mutFUS aggregates. IF images with FUS-WT-YFP are shown in Fig. S4A,F, S5D,E. (D) Immunofluorescent images of mOrange-tagged DNAJB12-FL and FLAG-tagged DNAJB14-FL showing co-localization of both chaperones within DJANGO structures. Cross correlation plots are shown together with the correlation coefficient (CC) between DNAJB14 and DNAJB12. See Fig. S4H for additional images. (E) Schematic representation of the DNAJB14 isoforms and mutants and their interactions with DNAJB12-FL and HSP70. (F,G) Co-IP between DNAJB12-FL (tagged with mOrange) and DNAJB14 isoforms and mutants, showing negligible interaction of DNAJB12-FL with DNAJB14-short, as well as reduced interaction with DNAJB14-ΔDUF. (G) Quantification of interaction was performed using Fiji, densitometry of mOrange bands were normalized to the corresponding FLAG bands (see Methods). Bars represent mean and std of 3 biological replicates (Additional replicates shown in Fig. S4J). (H) FUS-R521H-YFP Aggregation modulation scores in cells expressing DNAJB14-FL that underwent DNAJB12 knockdown (siDNAJB12+DNAJB14-FL) vs. cells treated with siControl (siControl+DNAJB14-FL). Bars represent mean and std error of N = 4 independent biological replicates. Dashed lines represent 95% CIs as in Fig. 2B. *** - p<0.003.

## DNAJB14 and DNAJB12 full-length isoforms form a complex

We subsequently examined the potential mutual role of DNAJB12 and DNAJB14 as a complex in the rescue of FUS-R521H-YFP aggregation. Each of these two chaperones contains a single transmembrane domain (TD, Fig. 1C,3A), and they were reported to be ER localized^27–29^. Indeed, DNAJB12-FL, DNAJB12-short and DNAJB14-FL were co-localized with an ER marker (Fig. S4C, see Methods), while DNAJB14-short that lacks the TD, did not (Fig. S4C). Immunofluorescence imaging of the four isoforms showed that DNAJB14-FL and DNAJB12-FL were able to generate unique nuclear structures (Fig. 3C,D, S4A,D-F,H), that were previously termed DJANGOs^26^. Neither DNAJB14-short nor DNAJB12-short could form the nuclear DJANGO structures (Fig. 3C and S4A,D-F). Immunofluorescence image analysis using a classifier trained to identify DJANGO structures (see Methods) confirmed that while DNAJB14-FL formed DJANGOs in about 10-14% of the cells, and DNAJB12-FL in about 7.5% of the cells, no DJANGOs were formed by DNAJB14-short or DNAJB12-short (Fig. S4G). In order to check whether DNAJB14-FL and DNAJB12-FL were co-localized in the cell, we tagged DNAJB12-FL with an mOrange fluorescent protein, and co-expressed it together with the FLAG tagged DNAJB14-FL. Immunofluorescence imaging showed that DNAJB12-FL and DNAJB14-FL co-localize to the same nuclear DJANGO structures (Fig. 3D, Fig. S4H). Moreover, these two chaperones were co-localized also in cells in which they both showed an ER localization pattern (Fig. S4I). Co-IP experiments confirmed that DNAJB12-FL and DNAJB14-FL physically interacted in cells, while DNAJB14-short interaction with DNAJB12-FL (Fig. 3F-G, S4J), and vice versa (S4K,L), were negligible.

## DNAJB14-mediated rescue of mutant FUS aggregation is HSP70 dependent

The fact that the full-length isoform of DNAJB14 substantially rescued FUS-R521H aggregation while the short isoform did not, hinted at the potential importance of the J-domain, and therefore of HSP70 interactions, in the rescue phenotype. Indeed, while DNAJB14-FL interacted with HSP70, DNAJB14-short did not, as shown by co-IP (Fig. S5A,B, 3E). To test the role of HSP70 interactions in the rescue phenotype, we generated a DNAJB14-FL version that was mutated in the HPD motif of the J-domain, the region where HSP70 is known to bind J-proteins^4^ (DNAJB14-HPDmut, Fig. 3A,E see Methods). This mutant showed negligible binding to HSP70 using co-IP (Fig. S5A,B), and this trend was consistent in cells expressing either FUS-WT-YFP or FUS-R521H-YFP (Fig. S5A,B). Immunofluorescence imaging of this mutant, as well as a second HPD mutant (DNAJB14-HPDmut2), demonstrated a substantially lower rate of DJANGO formation (Fig. 3C, Fig. S5C,D). Surprisingly, co-IP experiments indicated that DNAJB14-HPDmut1 showed enhanced binding to DNAJB12-FL (Fig. 3F,G, S4J). Finally, we tested the functional effects of the HPD mutations on FUS-R521H-YFP aggregation. Our data showed that these mutants lost their ability to provide significant rescue of FUS-R521H-YFP aggregation (Fig. 3B).

Thus, it seems that while HSP70 binding by DNAJB14-FL was not necessary for its interaction with DNAJB12-FL (Fig. 3E), the ability of DNAJB14 to interact with HSP70 was crucial for its function in alleviating FUS-R521H-YFP aggregation.

## DNAJB14 - DNAJB12 complex inter-dependence in the rescue of mutant FUS aggregation

We next examined the role of the interaction between DNAJB14-FL and DNAJB12-FL in the rescue of FUS-R521H-YFP aggregation. To that end, we used siRNA to knock down DNAJB12, to about half its endogenous levels (Fig. S6A). We then tested the functional effect of DNAJB14-FL ectopic expression on FUS-R521H-YFP aggregation in cells that were knocked down for DNAJB12 (see Methods). We found that knockdown of endogenous DNAJB12 reduced the ability of DNAJB14-FL to rescue FUS-R521H-YFP aggregation by about 50% (Fig. 3H). This indicated that the rescue of FUS-R521H-YFP aggregation provided by DNAJB14-FL is dependent on DNAJB12-FL, in addition to its interaction with HSP70.

## DNAJB14 DUF domain plays an important role in the DNAJB14-DNAJB12 complex formation and the rescue of mutant FUS aggregation

Our collective evidence thus far supported the notion that it is the complex of DNAJB14-DNAJB12 together with HSP70 that is important for the rescue of FUS-R521H-YFP aggregation. We next raised the possibility that the DUF domain (Domain of Unknown Function) of DNAJB14 might mediate the DNAJB14-DNAJB12 complex formation. To test this hypothesis, we generated a truncated DNAJB14 version lacking the DUF domain (termed DNAJB14-△ DUF, Fig. 3A,E see Methods). First, we examined the interaction of the DNAJB14-△DUF with HSP70 using co-IP, and found that the removal of the DUF domain reduced the interaction to about 50% (Fig. S6B,C). Furthermore, we found that, while DNAJB14-FL strongly interacted with DNAJB12-FL, DNAJB14-△ DUF interactions with DNAJB12-FL were substantially weaker, about 3 fold lower compared to DNAJB14-FL (Fig. 3F,G, S4J). This indicated that the DUF domain is involved in mediating DNAJB14-DNAJB12 complex formation. Finally, testing the functional effect of DNAJB14-△DUF on FUS-R521H-YFP aggregation phenotype revealed that the removal of the DUF domain severely compromised the rescue of FUS-R521H-YFP aggregation by DNAJB14 (Fig. 3B).

These results collectively establish that the DNAJB14-DNAJB12-HSP70 complex is essential for providing substantial rescue of mutFUS aggregation.

## DNAJB14 isoforms show differential modulation of mutant FUS aggregation in primary neurons

Finally, we wanted to examine the functional effects of the two naturally occurring DNAJB14 isoforms in primary neurons. To this end, we infected rat neuronal cultures with an AAV2 viral construct expressing either FUS-WT-YFP or FUS-R521H-YFP. Six days after infection, we observed a substantial amount of FUS aggregation within live neurons (Fig. 4A, S7A). The aggregation phenotype was different than that observed in HEK293T cells, namely, rather than a single large aggregate, neurons mostly exhibited several large cytoplasmic FUS-R521H-YFP aggregates along with multiple smaller FUS-R521H-YFP foci in the cytoplasm (Fig. 4A, Supplementary Movies S1,S2). The latter could also be seen, though less frequently, in neuronal projections (Fig. S7B). Nevertheless, in contrast to FUS-WT-YFP expressing neurons (Fig. 4B, Movie S3), neurons that showed FUS-R521H-YFP cytoplasmic aggregates were largely devoid of fluorescent signal in the area of the nucleus (Fig. 4A). We next tested the functional effects of co-expression of either DNAJB14-FL or DNAJB14-short on aggregate formation. Due to the complex phenotype of FUS-R521H-YFP aggregates, we subjected confocal microscopy images of live neurons to image analysis. We used a classifier trained to distinguish between nuclear fluorescent pattern (as in FUS-WT-YFP expressing neurons, Fig. 4B) and individual cell images displaying the complex FUS-R521H-YFP aggregate phenotype (see Methods). The classifier had a low background error rate, as shown by its application to FUS-WT-YFP infected neurons images (Fig. S7C,D, see Methods). We then quantified the percentage of aggregate containing neurons in FUS-R521H-YFP neuronal cultures co-infected with either DNAJB14-FL or DNAJB14-short, six days after viral infection. In total, we analyzed 4504 and 4381 FUS-R521H-YFP expressing neurons co-infected with DNAJB14-FL or DNAJB14-short, respectively. Strikingly, we observed that DNAJB14-FL expressing neurons showed significantly less FUS-R521H-YFP aggregation compared to DNAJB14-short expressing neurons (Fig. 4C,D, S7E,F). FUS-R521H-YFP neuronal cultures showed on average 26% less aggregate-containing cells when expressing DNAJB14-FL compared to DNAJB14-short (p=7.9e-7, 0.023, 5.7e-3, N=3, Fig. 4D, S7F, DNAJB14-FL and DNAJB14-short were titer-matched, see Methods). Therefore, our data demonstrated that the screen for mutFUS aggregation modulators performed in HEK293T human cells was able to identify chaperone modifiers that exhibited the same capability in modulating mutFUS aggregation in cultured primary neurons.

**Figure 4:**
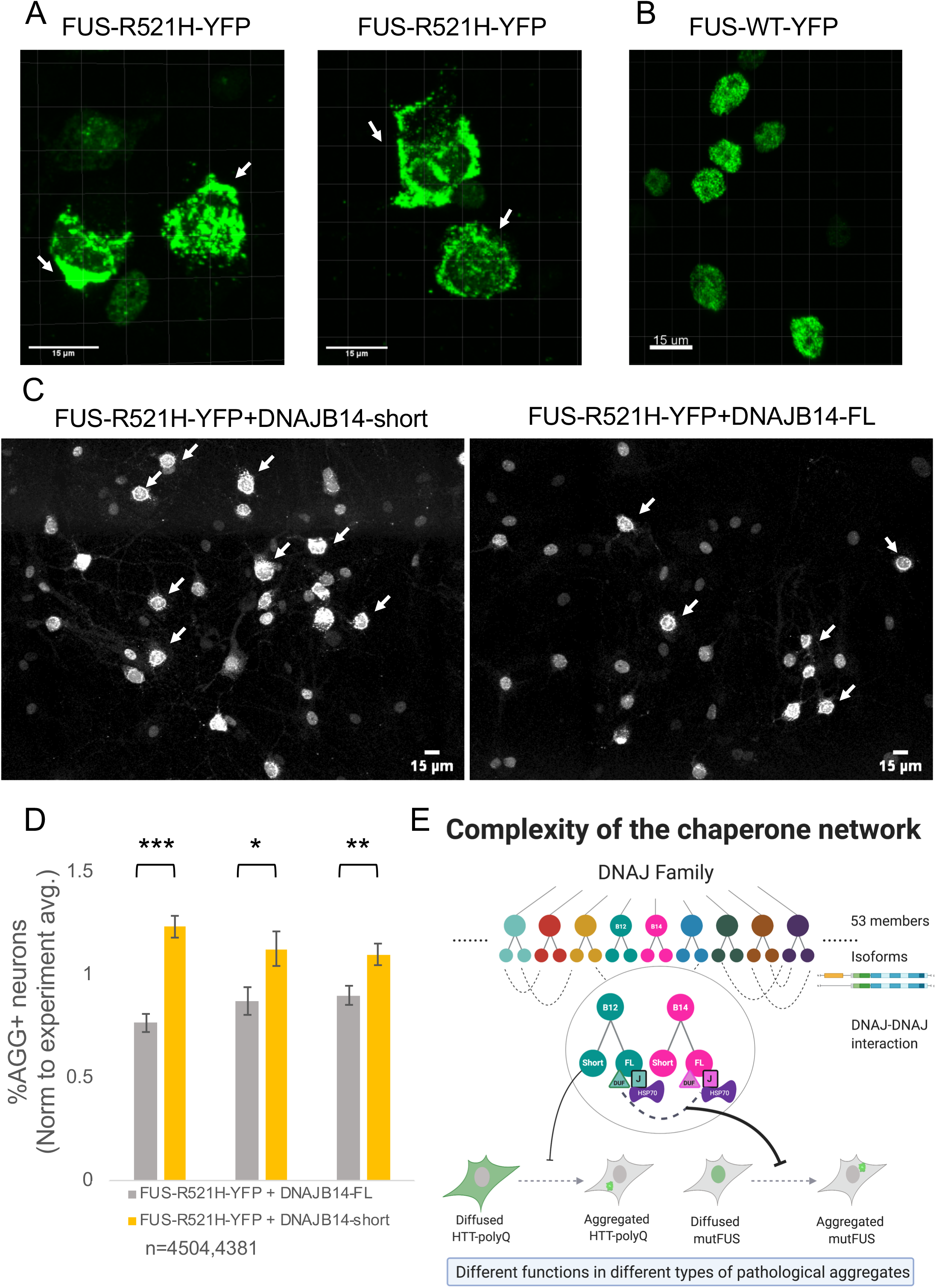
DNAJB14 isoforms show differential modulation of mutant FUS aggregation in primary neurons. (A) Confocal microscopy images of live neurons showing a complex FUS-R521H-YFP aggregation pattern. White arrows mark neurons with FUS-R521H-YFP aggregates. The left panel contains additional cells with a non aggregated FUS-R521H-YFP expression pattern. Visualized using Imaris. (B) Confocal microscopy images of live neurons expressing FUS-WT-YFP. Visualized using Imaris. (C) Larger fields of neuronal cultures co-infected with FUS-R521H-YFP and either DNAJB14-short (left panel) or DNAJB14-FL (right panel). Shown are maximum-projection of confocal fluorescent images (using Fiji). Additional fields are in Fig. S7E. White arrows mark neurons with mutFUS aggregates. (D) Image analysis of thousands of live neurons showed significantly lower FUS-R521H-YFP aggregation in DNAJB14-FL expressing neurons compared to DNAJB14-short. Neuronal cultures were co-infected with FUS-R521H-YFP and either DNAJB14-FL or DNAJB14-short containing viruses with equal titers at day 5 of culture, and imaged six days later using confocal microscopy. Graphs show mean and std error of the aggregation fraction, normalized to the average aggregation fraction measured overall in each experiment, in three independent biological replicate experiments. Experiment average aggregation fraction was calculated including all fields of both DNAJB14-FL and DNAJB14-short samples. Experiments included 12, 24 and 37 images from each of DNAJB14-FL or DNAJB14-short samples, containing a total of 884 and 1081, 1339, or 1069, 2281 and 2231 neurons for DNAJB14-FL or -short respectively. All experiments showed significant differences (difference were 37%, 22% and 18%, p=7.9e-7, 0.023 and 5.7e-3 for the three replicates, calculated using student t-test, see Methods). Non normalized data presented in Fig. S7F. (E) Naturally-occurring DNAJ isoforms and their interactions show differential functions with respect to different aggregate types, revealing inter-family functional diversification, thereby increasing the complexity of the chaperone network.

## Discussion

Here we identified a complex of DNAJB14-FL and DNAJB12-FL, which interact with HSP70 to significantly protect cells from mutFUS aggregation (Fig. 4E). Importantly, naturally occurring short isoforms of both DNAJB12 and DNAJB14, which could not participate in a complex or interact with HSP70, were unable to significantly rescue mutFUS aggregation. Conversely, DNAJB12-short showed HTT-polyQ aggregation protection. Thus, our work revealed that naturally-occurring DNAJ isoforms show different functional diversification towards different types of aggregates.

Although WT FUS is predominantly nuclear, it also exhibits nucleocytoplasmic shuttling^30^. A class of ALS mutations in FUS, including R521H and R518K examined here, lead to FUS aggregation outside of the nucleus (Fig. 2A,4A). Notably, FUS-R521H showed diffused nuclear localization in 26-40% of the cells (based on image analysis, Fig. S4A, see Methods), but aggregates were rarely nuclear (1.5-2.6% of the cells, based on image analysis, see Methods). The nuclear import factor Kap-beta2 was shown to inhibit FUS-R521H aggregation in cells^31^, supporting the notion that keeping mutFUS nuclear prevents its aggregation. DNAJB12 and DNAJB14 were previously shown to have roles in ERAD on the one hand^27–29^, and form the nuclear DJANGO structures^26^, whose function is currently unknown, on the other hand. Using image analysis, we found that DNAJB14-FL DJANGOs were present in about 10-14% of our cells (Fig. S4G). Our data demonstrated that the ability to form DJANGOs was correlated with the ability to rescue mutFUS aggregation. An intriguing hypothesis is that these structures participate in regulating the shuttling of mutFUS, thereby aiding its solubility in the nucleus, however the potential involvement of DJANGOs in mutFUS aggregation modulation remains for future exploration.

Interestingly, we found that while the rescue of mutFUS aggregation was HSP70 dependent, rescue of HTT-polyQ by DNAJB12-short was not only HSP70 independent, but the DNAJB12-FL isoform that interacts with HSP70 aggravated HTT-polyQ aggregation (Fig. 1B). Interestingly, in the case of the previously identified HTT-polyQ aggregation modulators DNAJB6 and DNAJB8^12^, aggregation protection was shown to be, at least in part, HSP70 independent^12^. It therefore seems that HTT-polyQ aggregation protection can be performed by several DNAJ proteins in an HSP70-independant manner. In contrast, as we found here, mutFUS aggregation alleviation required HSP70 in the context of the DNAJB14-DNAJB12 complex. These findings highlight different chaperoning requirements from these different types of pathological aggregates.

The chaperone network in general, and the HSP40 (DNAJ) family in particular, has undergone a major expansion in evolution, and the human genome contains 53 different DNAJ proteins^4^. Our work reveals that the interplay between naturally-occurring isoforms of DNAJs further increases the complexity of the chaperone network, by generating functional diversification towards different types of pathological aggregates (Fig. 4E). Therefore, modulation of protein homeostasis in general, and of protein aggregation in particular, can potentially be performed by isoform switching of DNAJs, in yet to be identified conditions. This further increases the complexity of the chaperone network as we view it today (Fig. 4E), and in addition to overexpression of specific chaperone isoforms as shown here, has the potential to further serve as a point for therapeutic intervention.

## Methods

### Plasmid preparation

HTT134Q-GFP and HTT17Q-GFP containing plasmid were a gift from Prof. Noam Ziv, and were cloned into the pTREX backbone. A FUS-YFP construct was purchased from Addgene ^19^, and then cloned into the pTREX backbone using PCR and gateway cloning. Subsequently, the Quikchange 2 Site-Directed Mutagenesis Kit (Agilent Technologies 210518) was used to introduce the two mutations: Arginine (R) to Histidine (H) in position 521, and Arginine (R) to Lysine (K) in position 518 of the protein. C-terminal FLAG-tagged library of 65 chaperones was prepared from the ORFeome library (purchased from GE Healthcare), and transferred from pDONR223 into pcDNA3.1-ccdb-FLAG using gateway cloning. Plasmids expressing mOrange-tagged proteins were prepared using gateway cloning into a pcDNA3.1-ccdb-mOrange backbone, which was generates in house. In addition, four isoforms of DNAJB14 were generated (all primers are listed in Table S2): DNJB14-short lacking the J-domain, TM and DUF domain (Fig. 3A) was generate from the full-length isoform using PCR and gateway cloning; two mutations in the HPD domain Histidine (H) to Glutamine (Q) in position 136 of the protein, introduced using the Quikchange 2 Site-Directed Mutagenesis Kit; DNAJB14△DUF truncation was performed using PCR from the full-length isoform and gateway cloning (see Fig. 3A). DNAJB14 and DNAJB12 isoforms accession numbers: DNAJB14-FL - ENST00000442697 (NM_001031723/uc003hvl), DNAJB14 short - uc003hvm.4 (similar to ENST00000469942.1), DNAJB12-FL - ENST00000394903 (NM_017626), DNAJB12-short - ENST00000461919 (KJ902687).

The pAAV-FUS-R521H-YFP, pAAV-FUS-WT-YFP, pAAV-DNAJB14-FL and pAAV-DNAJB14-short plasmids were constructed using Gateway cloning into pAAV-CAG-dest. pAAV-CAG-dest was a gift from Prof. Yitzhak Kehat. An ER marker plasmid, mCherry-ER-3, containing the CALR signal peptide region fused to mCherry, was a gift from Michael Davidson (Addgene plasmid #55041).

### Cell culture and Transfection

HEK293T cells were grown in standard DMEM supplemented with 10% FBS, 1% Penicillin-Streptomycin in 37°C, 5% CO2 incubator. For the FUS screen transfections, HEK293T cells (3e5 cells per well) were seeded on 6well plates and transfected using PEI transfection reagent (Thermo Fisher Scientific) one day after seeding, according to the manufacturer instruction, with 1250 ngr of FUS DNA and 1250 ngr of chaperone DNA. For HTT screen transfections, cells were seeded at 25,000 cells per well on 96 well plates one day before transfection, and transfected using PEI with 200 ngr of HTT DNA and 200 ngr of chaperone DNA. Cells were assayed 48 hours following the transfection.

### PulSA method

At 48h after transfection, cells were trypsinized, resuspended in media with serum, placed on ice, and DAPI regent was added (5μM) in order to assay cell viability. Cells were then taken to Flow cytometry in order to differentiate between subpopulations with diffused cellular fluorescence and those with protein aggregates, as in Ramdzan et al. ^21^, and as illustrated in Fig. 1A. To obtain Aggregation modulation scores, cells of appropriate size were considered during the FACS analysis, and non-viable cells (positive for DAPI) were excluded. We then quantified the percentage of cells that showed a fluorescent aggregate pattern (AGG+), and compared these percentages to cells co-transfected with DsRed as a control. Aggregation modulation scores for each chaperone were calculated as Log2(%AGG+ (chaperone)/%AGG+(DsRed)). Each experiment was performed in 4 independent biological replicates for 66 chaperones. Confidence intervals (95%, appear as dashed lines in Fig. 1B, 2B etc.) were calculated according to the variations between different DsRed co-transfected cells replicates, such that they represent twice the standard deviation of Aggregation modulation scores calculated between different replicates of DsRed performed on the same day. In order to find an appropriate baseline control for the screen, we tested aggregate rates in cells co-transfected with each of two unrelated proteins: Renilla luciferase, or DsRed, while also comparing to cells transfected with aggregated protein alone. As co-expression of DsRed gave lower levels of basal aggregation rates (Fig. S1D,E, S3C), we chose the more restrictive baseline and used DsRed as a control.

### Dose-dependent aggregation modulation scores

To check the effect of different doses of chaperones on aggregation modulation, cells were transfected one day after seeding (3e5 cells per well in a 6well plate) in three doses: 2000ngr of FUS DNA plus 500ngr of chaperone DNA, 1250ngr of FUS DNA plus 1250ngr of chaperone DNA, and 700ngr of FUS DNA plus 1800ngr of chaperone DNA (Fig. 2C). Cells were assayed 48 hours following transfection by PulSA as above. Each combination was compared to a respective combination of FUS DNA plus DsRed DNA to obtain the Aggregation modulation score.

### Immunofluorescence staining

Cells were cultured on cover-slips and transfected as above. Two days post transfection, cells were fixed in 4% paraformaldehyde in PBS, permeabilized with 0.5% Triton-x100 in IF buffer (5% FCS, 2% BSA in PBS), then immunostained by primary anti-FLAG antibody (Sigma-Aldrich, F1804) followed by secondary AlexaFluor 647 Donkey Anti-Mouse (Jackson) antibody to image FLAG-tagged chaperones. The far-red secondary antibody was used to eliminate fluorescence overlap between the YFP/GFP-tagged aggregates and the chaperones. The cells were additionally stained with DAPI to mark nuclei.

### Image analysis

A Fiji script was automatically run on all images identifying cells containing both nuclear marker (DAPI) and FLAG tagged chaperone staining (as above). This generated a binary image that contained cells above a fixed threshold for each channel. After size filtering and signal thresholding cells were counted.

DJANGO structure identification was done using the Trainable Weka Segmentation plugin, followed by summation of the signal and thresholding.

Aggregates were detected using the Trainable Weka Segmentation plugin, and so were cells with diffused FUS-R521H-YFP fluorescence. Aggregate localization was determined using summation of the identified aggregate signals either inside the nucleus (which were modeled by the DAPI channel) or outside (using a larger definition of a fixed area around the nucleus, and subtraction of any aggregates signal that was identified within this area from the total aggregate area calculated per cell).

### Co-immunoprecipitation

Co-immunoprecipitation (co-IP) lysis buffer was prepared as follows: 50mM HEPES pH 7.9, 0.5% Triton X-100, 5% glycerol, 150mM NaCl and 2 mM EDTA and Protease inhibitors (Sigma–Aldrich P5726). Wash buffer was prepared as follows: 50mM HEPES pH 7.9, 1% Triton X-100, 5% glycerol, 150mM NaCl and 2 mM EDTA. Cells were transfected as above. At 48h post transfection, cells were placed on ice, washed with ice cold PBS, lysis buffer was added and cells were collected by scraping. Lysates were centrifuged (8200g for 10 minutes at 4°C), and supernatant was transferred to new tubes. Samples were IP-ed using 10μl of EZview Red ANTI-FLAG M2 Affinity Gel beads (Sigma-Aldrich, F2426), which were added to a sample volume containing equal number of 260 nm OD units. Additional co-IP buffer was added to reach a 600μl volume for each sample and samples with beads were slowly rotated 3h at 4°C. Beads were then washed 7 times with 1ml of wash buffer. Following the last wash, the beads were resuspended in equal volumes of wash buffer, protein sample buffer was added, and the samples were boiled at 100°C for 5 minutes. Equal volumes were then run on an 8% or a 12.5% gel, and immunoblotted using anti-FLAG (Sigma-Aldrich, F1804), anti-HSC70/HSP70 (Enzo Life Sciences, N27F34), or anti-RFP (MBL, 598) antibody, which was used to detect mOrange tagged proteins. Immunoblots densitometry was quantified using Fiji, where interactor levels (HSP70 or proteins tagged with mOrange) were normalized to the pulled-down protein levels (FLAG).

### Small interfering RNA

Cells were seeded at 150,000 cells per 6well plate one day before transfection. siRNA transfections were performed using RNAiMAX lipofactamine reagent (Thermo Fisher Scientific), according to manufacturer’s protocol, with siRNA concentration of 15nM. One day after transfection of the siDNAJB12 or siControl (Dharmacon Smartpool, 001810-10-05 or 020585-01-0005), cells were co-transfected with 1250ngr of FUS-R521H and 1250ngr of DNAJB14-FL using PEI. Cells were analyzed 48h later.

### Virus production

All viruses were packaged with the AAV-PHP.B capsid using an optimized protocol by Challis et al. ^32^ for producing AAV viruses from HEK293T cells grown on six 10cm plates. All viruses were then tittered using qPCR in the same assay and dosage consistency was kept between pAAV-FUS-R521H-YFP and pAAV-FUS-WT-YFP (infections made at the same titer), and between pAAV-DNAJB14-FL and pAAV-DNAJB14-short (infections made at the same titer).

### Neuronal cultures and infections

Isolation and culturing of rat primary hippocampal neurons were done as previously described ^33^. Briefly, P0 neonates, from timed-pregnant white coat female Sprague Dawley female rats (Charles River; SAS SD, strain code 400), were euthanized and their hippocampi dissected and dissociated. Then, 150K cells were plated in 24-well flat bottom plates on coverslips (12 mm, thickness ~0.1 mm) coated with Poly-D-Lysine (Sigma-Aldrich P7886-50MGP) in Minimum essential medium (MEM), glutamine-free based Neuronal growth medium (5% fetal bovine serum, 2% B27, 1% Glutamax (100×, Invitrogen), 2 % 1 mol/L d-glucose, 0.1% μL serum extender (BD Biosciences, Cat. No. 355006)). Cultures were incubated at 37°C and 5% CO_2_. At 5 days after plating, media was changed and supplemented with cytosine arabinosine (Ara-C, Sigma, C6645) at a final concentration of 4 μM, to reduce the growth of glia in the cultures. Cultures were infected the same day with pAAV-FUS-R521H-YFP or pAAV-FUS-WT-YFP (at the same titer), together with either pAAV-DNAJB14-FL or pAAV-DNAJB14-short (at the same titer). After that media was changed every two days, and cells were imaged six days after infection.

### Neurons image analysis

For microscopy-analysis imaging, co-infection of pAAV-FUS-R521H-YFP or pAAV-FUS-WT-YFP with equal dosages of either pAAV-DNAJB14-FL or pAAV-DNAJB14-short virions occurred at the 5th day post primary cell culture isolation, imaging took place at the 6th post infection.

Images were acquired with a Zeiss LSM 880 confocal microscope using water dipping Plan-Apochromat ×20 objective, (working distance, 1.8 mm). The acquired images were processed and analyzed using a Fiji script. In short, out of each confocal microscopy image, 15 slices were successively chosen along the z-axis and stacked to form one image. An equal number of regions of interest (651×651 μm in size) were randomly chosen as input for the script. Then, total cell numbers per processed image were identified based on intensity threshold (>240), and images were matched between DNAJB14-FL and DNAJB14-short sets to have approximately the same number of cells. Subsequently, separation of cells was performed based on watershed processing, cell filtering was based on size (>50 μm2), and cellular aggregate detection was performed using Trainable Weka Segmentation plugin, summing aggregate area per cell, where a total area >5 μm were considered aggregates. Finally, a Python script was used to extract output excel files.

Each biological replicate, which included either pAAV-FUS-R521H-YFP or pAAV-FUS-WT-YFP (at equal titers), each co-infected together with pAAV-DNAJB14-FL or pAAV-DNAJB14-short (at equal titers) was imaged to obtain thousands of neurons. For pAAV-FUS-R521H-YFP we obtained 12, 24 and 37 images from each of DNAJB14-FL or DNAJB14-short samples (containing a total of 884 and 1081, 1339 and 1069, 2281 and 2231 neurons for DNAJB14-FL and -short respectively). For pAAV-FUS-WT-YFP we obtained 16 images for each of DNAJB14-FL and DNAJB14-short samples (containing 741 and 883, 659 and 689, 1131 and 1208 neurons for DNAJB14-FL and -short respectively). These images were subjected to image analysis as described above. The number of aggregate-containing cells was divided by the total number of fluorescent cells detected by the classifier to obtain a fraction of aggregate containing neuron for each image. Then, average and std error for the fraction of aggregate-containing neurons was calculated for DNAJB14-FL and -short in each experiment, and a two-tailed t-test p-value was calculated for the difference between the fraction of aggregate-containing neurons calculated for DNAJB14-FL and -short in each experiment. The image analysis classifier had a low background error rate, demonstrated by its application to images taken from FUS-WT-YFP expressing neurons, and no consistent difference was observed between DNAJB14-FL and DNAJB14-short FUS-WT-YFP co-expressing neurons. Overall 1.3% and 2.3% aggregation rate was identified for DNAJB14-FL and DNAJB14-short respectively in FUS-WT-YFP expressing neurons images (Fig. S7C).

## Acknowledgements

We thank the ISF and the Prince Center for Neurodegenerative disorders for their support in funding this project. We thank Herman Wolosker for critical reading of the manuscript.

## Author contributions

The project was conceived by RS. RS, KR and AY designed all experiments. KR and AY performed all experiments. LK performed all image analysis. SB extracted primary neurons and RH helped with viral production. RS wrote the manuscript together with KR and AY.

## Competing interests

The authors declare no competing interests.

**Supplementary Information** is available for this paper.

## Data Availability

The datasets generated during the current study are included in this published article and its supplementary information files. Raw data are available from the corresponding author upon request.

## Notes

### Competing Interest Statement

The authors have declared no competing interest.

